# Insights into the evolution of extracellular leucine-rich repeats in metazoans with special reference to Toll-like receptor 4

**DOI:** 10.1101/269241

**Authors:** Dipanjana Dhar, Debayan Dey, Soumalee Basu

## Abstract

The importance of the widely spread leucine-rich repeat (LRR) motif has been studied considering TLRs, the LRR-containing protein involved in animal immune response. The protein connects intracellular signalling with a chain of molecular interaction through the presence of LRRs in the ectodomain and TIR in the endodomain. Domain analyses with human TLR1-9 reported ectodomain with tandem repeats, transmembrane domain and TIR domain. The repeat number varied across members of TLRs and remains characteristic to a particular member. Analysis of gene structure revealed absence of codon interruption with TLR3 and TLR4 as exceptions. Extensive study with TLR4 from metazoans confirmed the presence of 23 LRRs in tandem. Distinct clade formation using coding and amino acid sequence of individual repeats illustrated independent evolution. Although ectodomain and endodomain exhibited differential selection pressure, however, within the ectodomain, the individual repeats displayed positive, negative and neutral selection pressure depending on their structural and functional significance.

## 1 Introduction

Leucine-rich repeat (LRR) containing domains are abundant in a diverse array of organisms, ranging from bacteria and archaea to eukaryotes. A number of databases such as SMART, Pfam, PROSITE and InterPro reports the presence of huge number of proteins with this highly repeating consensus motif (Sonnhammer et al., 1997; Sigrist et al., 2010; Burge et al., 2012; Letunic et al., 2012). Tandem arrays of characteristic LRR motifs consisting of approximately 20-30 amino acids and the repeat numbers ranging from 2 to 62 have been found in the primary structure of a large number of proteins (Kajava, 1998; Kobe and Kajava, 2001; Matsushima et al., 2010) encompassing a wide range of functions. Each repeat is segmented into a highly conserved segment (HCS) and a variable segment (VS) (Kajava et al., 1995; Ohyanagi and Matsushima, 1997). The HCS usually consists of either the 11-residue sequence LxxLxLxxNxL or the 12-residue sequence LxxLxLxxCxxL, where L is Leu, Ile, Val, or Phe; N is Asn, Thr, Ser, or Cys; and C is Cys, Ser, or Asn (Kajava, 1998; Kobe and Kajava, 2001). Although these substitutions often preserve hydrophobicity or polarity, it is possible for the first or the last leucine to be replaced by a relatively hydrophilic residue (Matsushima et al., 2007). Based on different length and consensus sequence of the VS, LRRs can be classified into 8 classes: “Typical”, “SDS22-like”, “Bacterial”, “Plant Specific (PS)”, “TpLRR (*Treponema pallidum* LRR)”, “RI-like (Ribonuclease inhibitor-like)”, “IRREKO” and “Cysteine-containing (CC)” (Kajava, 1998; Kobe and Kajava, 2001; Matsushima et al., 2010). LRR-containing proteins form a characteristic solenoid to provide a structurally invariant scaffold for protein-protein interaction. The concave side of the solenoid is defined by a series of parallel β-sheet, each constituted by HCS of an LRR motif whereas the convex side is comprised of α-helices, 3_10_ helices, polyproline II helices, β-turns and even short β-strands (Kobe and Deisenhofer, 1994; Kajava et al., 1995; Bella et al., 2008; Botos et al., 2011).

One of the prime functions of LRR involves innate immunity. For example, in organisms like invertebrates (e.g. sea urchin) and Cephalochordates (e.g. amphioxus), which lack adaptive immunity, vast repertoires of LRR proteins are present (Huang et al., 2008). Beyond innate immunity, LRRs being versatile binding motifs exhibit extensive functional significance in a variety of cellular processes like apoptosis, autophagy, cell polarization, neuronal development, nuclear mRNA transport, regulation of gene expression and ubiquitin-related processes (Kobe and Deisenhofer, 1994; Kobe and Kajava, 2001; Matsushima et al., 2005; Matsushima et al., 2005a; Bella et al., 2008; de Wit et al., 2008; Ng and Xavier, 2011). On the basis of domain organization of LRR in a protein, they can be classified into two major types: in tandem repeats [e.g. Ribonuclease inhibitor (RI), Polygalacturonase-inhibiting protein (PGIP) and Yersinia outer protein M (YopM)] or in association with other domains like Toll/interleukin-1 receptor (TIR), Gelsolin, and NACHT [e.g. Toll-like receptor (TLR), Protein Flightless-1 homolog, NACHT LRR and PYD domains-containing protein 6]. In this study, we have focused on the animal protein TLR, involved in innate immune response.

Toll-like receptors, initially discovered as Toll in fruit fly, consists of ten transmembrane glycoproteins in humans (TLR1-10) which recognizes a wide range of pathogen-associated molecular patterns (PAMPs) from bacteria, virus, fungi as well as some host molecules (Takeda et al., 2003). The domain organization is common across all the TLRs and they have one extracellular domain composed of several LRRs, a helical transmembrane domain and an intracellular TIR domain (Medzhitov, 2007). The extracellular domain recognizes various types of PAMPs. For example, lipopeptides or lipoproteins are recognized by TLR2 forming a heterodimer with TLR1 or TLR6, lipopolysaccharides (LPS) are recognized by TLR4 in association with myeloid differentiation factor 2 (MD-2), bacterial flagellin by TLR5 and viral double stranded and single stranded RNAs are identified by TLR3 and TLR7, TLR8, respectively (Akira, 2003; West et al., 2006). In terms of recognition, TLR10 remains an orphan receptor with no characteristic agonists and lacks a homologue in mouse (Hasan et al., 2005). The TIR domains are associated with intracellular signalling pathways that are regulated by various adapters (e.g. MyD88, TIRAP/Mal, TRIF and TRAM). Differential utilization of these TIR domain recruiting adapters provides selection of TLR mediated signalling pathways. Some TLRs are cell surface bound (TLR1, TLR2, TLR4, TLR5, TLR6 and TLR10) whereas some are sequestered in endosomal compartments (TLR3, TLR7, TLR8 and TLR9). Consequently, the ectodomain portion are either extracellular or face the lumen of an intracellular compartment where they encounter the pathogen derivatives.

The ectodomain part of the TLR varies notably in context to the number of LRR motifs and the amino acid residues comprising them. These variations help in the recognition of a broad range of ligands thereby activating the signalling cascade through the TIR domain via a defined set of adapter molecules for specific TLRs. It may be mentioned here that the number of signalling pathways and adapter proteins utilized for the same are relatively smaller than the vast spectrum of the recognition machinery of TLRs. The intriguing question that arises is whether the residues (in HCS or VS) of the LRR motif or the number of repeats contribute independently or in combination to play a pivotal role in recognition and initiation of a streamlined immune response. Thus, the ectodomain has to evolve at a higher rate, to protect the host from the repertoire of constantly evolving pathogens, by presumably maintaining co-evolutionary dynamics. There have been numerous views on the evolution of TLRs. Initially, it was thought that evolutionary conservation and strong functional constraint exist on TLRs (Roach et al., 2005). Alternatively, it has been suggested that functional heterogeneity of TLRs arise from occurrence of gene duplication, gene loss and gene conversion (Hughes and Piontkivska, 2008; Roach et al., 2013). Moreover, variation in the ectodomain region of TLRs has been found to be high compared to the TIR domain (Mikami et al., 2012). Thus, we examined evolutionary relationships in TLRs (taking whole protein, ectodomain and endodomain part individually) by generating phylogenetic trees. In addition, we have studied differential selection pressure on distinctive regions of TLRs specifically on the individual LRRs enriching the ectodomain part.

## 2 Materials and methods

### 2.1 Sequence Data

The amino acid sequences of human TLR1-9 were collected from the National Center for Biotechnology Information (NCBI) database (https:www.ncbi.nlm.nih.gov/protein/) on October, 2017 and the genes for the same were retrieved using the Ensembl Genome Browser v90 (Aken et al., 2017). The following sequences from humans with their respective GI number were considered: TLR1 [49065550], TLR2 [21708105], TLR3 [86161330], TLR4 [6175873], TLR5 [80478954], TLR6 [5006248], TLR7 [74096726], TLR8 [37181712] and TLR9 [74054321]. The sequences were retrieved through extensive keyword match and BLAST (Altschul et al., 1990) searches. Only complete sequences were considered. Each sequence, corresponding to a distinct TLR, was matched for a Uniprot ID and only those having an ID were selected. It may be added that precursor/partial sequences were not considered in the study. Amino acid and gene sequences of TLR4 were obtained for six other metazoans (monkey, rat, mouse, sheep, cow and horse) from NCBI and Ensembl Genome Browser v90 respectively.

### 2.2 Determination of Domain Architecture of TLRs

The protein sequences were segmented into ectodomain (containing LRR motifs and transmembrane region) and endodomain (consisting of TIR). This division was done based on the results from NCBI Conserved Domain Database (CDD), SMART and UniProt databases. Multiple sequence alignment was performed separately on ectodomain and endodomain of the nine TLRs, using ClustalW (Thompson et al., 1994) and T-Coffee (M-Coffee) (Notredame et al., 2000; Poirot et al., 2003) with the set of default parameters for MSA. The results were saved in PHYLIP output format, for phylogenetic tree construction.

### 2.3 Study with the gene sequences

The coding sequences and the exon-intron structures of seven organisms for each TLR (TLR1-9) were obtained from the annotations associated with sequences in the Ensembl Genome Browser v90.The gene sequences retrieved from Ensembl were selected in a manner that is provided with RefSeq and Uniprot ID. All the sequences selected belong to transcript level 1 (TSL1).

### 2.4 LRR identification within TLR4

The protein sequences of TLR4 from nine different organisms were submitted to the SMART and Pfam database in order to calculate the number of LRRs. Following disparities in the results from both the databases, LRRs were counted manually and re-checked to ensure that substitutions or conserved mutations were not overlooked. Multiple sequence alignment was performed on TLR4 protein from nine organisms with the following GI numbers: human [6175873], monkey [194068441], horse [9717253], cow [111414451], goat [334812887], sheep [198281858], buffalo [324121078], mouse [148699131] and Norway rat [149059567]. LRR motifs were marked on the MSA after thorough evaluation based on the usual conserved pattern i.e. L x x L x L x x N/C x L.

### 2.5 Phylogenetic Tree Analysis

Phylogenetic trees showing evolutionary relationships among the human TLRs were constructed using the protein sequence parsimony method (Protpars) in the PHYLIP (phylogeny inference package) program (Felsenstein, 1989). Three types of trees were made considering the entire TLR protein, ectodomain and endodomain separately. Besides, phylogenetic trees were established using the TLR4 whole protein and 23 individual LRRs of the ectodomain region across seven organisms as the genes for buffalo and goat were unavailable in the Ensembl Genome Browser v90. A total of 1000 bootstrap values were used to generate bootstrap replicates utilizing the seqboot program of the PHYLIP package. On reaching convergence, all the trees were explored and a consense tree was initiated using the consense program of the package. FigTree v1.4.3 (http://tree.bio.ed.ac.uk/software/figtree/) and iTOL v4 (Letunic and Bork, 2016) were used collectively for both initial visualization and generation of figures of the resulting phylogenetic trees.

Similarly, phylogenetic trees using bootstrapping were constructed for the coding sequences of TLR4 among seven organisms. In addition, a maximum parsimony tree was generated with exon sequences corresponding to 23 LRR regions for each organism using dnapars program in the PHYLIP package.

### 2.6 Estimation of K_a_/K_s_ ratio

K_a_/K_s_ or dN/dS indicate the ratio of non-synonymous to synonymous substitution rate. Under neutrality, the ratio is not expected to deviate significantly from 1 (ω = 1) whereas notable changes in the ratio may be interpreted in case of positive selection (ω>>1) or negative selection (ω<<1). The ratio was determined for the coding sequence of 23 LRRs individually taken from TLR4 of seven organisms, using an online available server known as K_a_/K_s_ calculation tool provided by Computational Biology Unit of Bergen Centre for Computational Science (http://services.cbu.uib.no/tools/kaks; Liberles, 2001). Additionally, the ratio was calculated for TLR1-9 of human and monkey using PAL2NAL (Suyama et al., 2006) considering three cases-ectodomain, endodomain and the whole protein.

## 3 Results & Discussion

### 3.1 Domain Organization in TLRs

All mammalian TLR1-9 are known to consist of unique number of LRRs along with a conserved TIR domain. The SMART and PROSITE databases were consulted to find out the number of LRRs and the position of the TIR domain. The number and position of LRRs differed significantly in these databases. Also, as evident from Figure 1, specific TLRs, namely TLR7, TLR8 and TLR9, are devoid of the TIR domain according to the SMART database, declaring it as an unknown region. We therefore consulted other databases like NCBI and UniProt, the cumulative result of which is tabulated in Table 1. Finally, the amino acid sequences of human TLR1-9 were segmented into two portions, one containing the ectodomain part (comprising the LRRs and TM region) and the other being the conserved endodomain part, bearing the TIR domain based on the above analysis.

**Table 1.** Number of LRRs in nine human TLRs predicted by four databases.

|  | *Number of LRRs predicted by* |  |  |  |
| --- | --- | --- | --- | --- |
|  | *SMART* | *PROSITE* | *NCBI* | *UNIPROT* |
| TLR1 | 4 | 10 | 19 | 19 |
| TLR2 | 7 | 11 | 19 | 19 |
| TLR3 | 16 | 19 | 22 | 22 |
| TLR4 | 11 | 11 | 18 | 18 |
| TLR5 | 9 | 12 | 22 | 22 |
| TLR6 | 5 | 9 | 19 | 19 |
| TLR7 | 12 | 18 | 27 | 27 |
| TLR8 | 15 | 20 | 23 | 23 |
| TLR9 | 18 | 17 | 26 | 26 |

**Figure 1.**
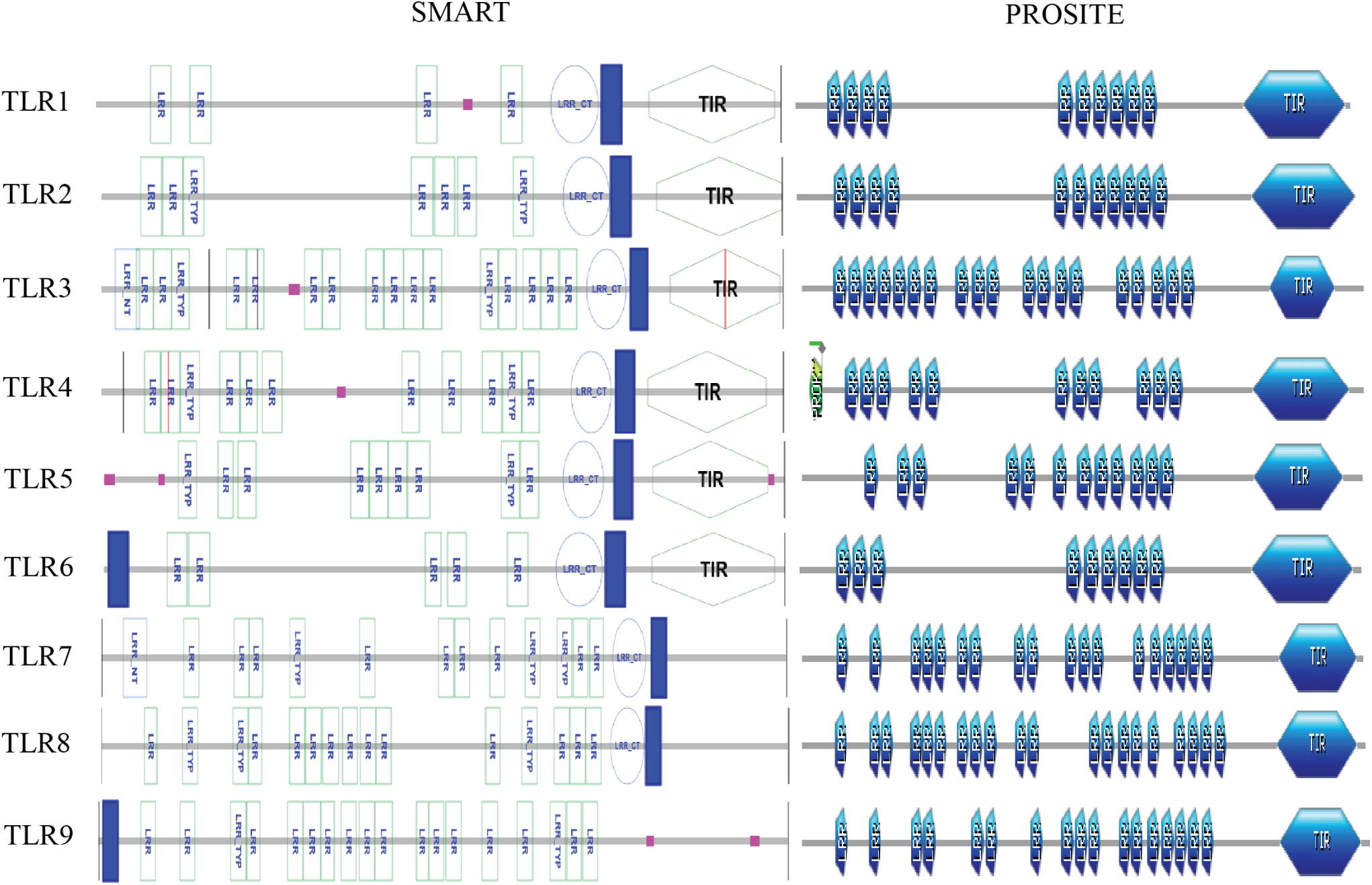
Domain architecture of human TLR1-9 as reported by SMART and PROSITE.

### 3.2 Evaluating the genes

The exon number, its length and phase of TLR1-9 from seven organisms are summarized in Figure 2. Phases of exons are expressed in pairs corresponding to the phases for the 5’ and the 3’ end of the exon. For the 5’ end, phase zero means that the exon begins with a complete codon, phase one means that the first two bases at the 5’ end of the exon constitute the second and third bases of an interrupted codon. Similarly, phase two means that the first base of the exon consists of the last base of an interrupted codon (Sharp, 1981). The phase at the 3’ end of the exon is determined by the phase of the following intron. On the other hand, introns that interrupt the reading frame and lie between codons are known as ‘phase zero’ introns, those that interrupt the codons between the first and second base are known as ‘phase one’ introns, and those interrupting between the second and third base are known as ‘phase two introns’.

**Figure 2.**
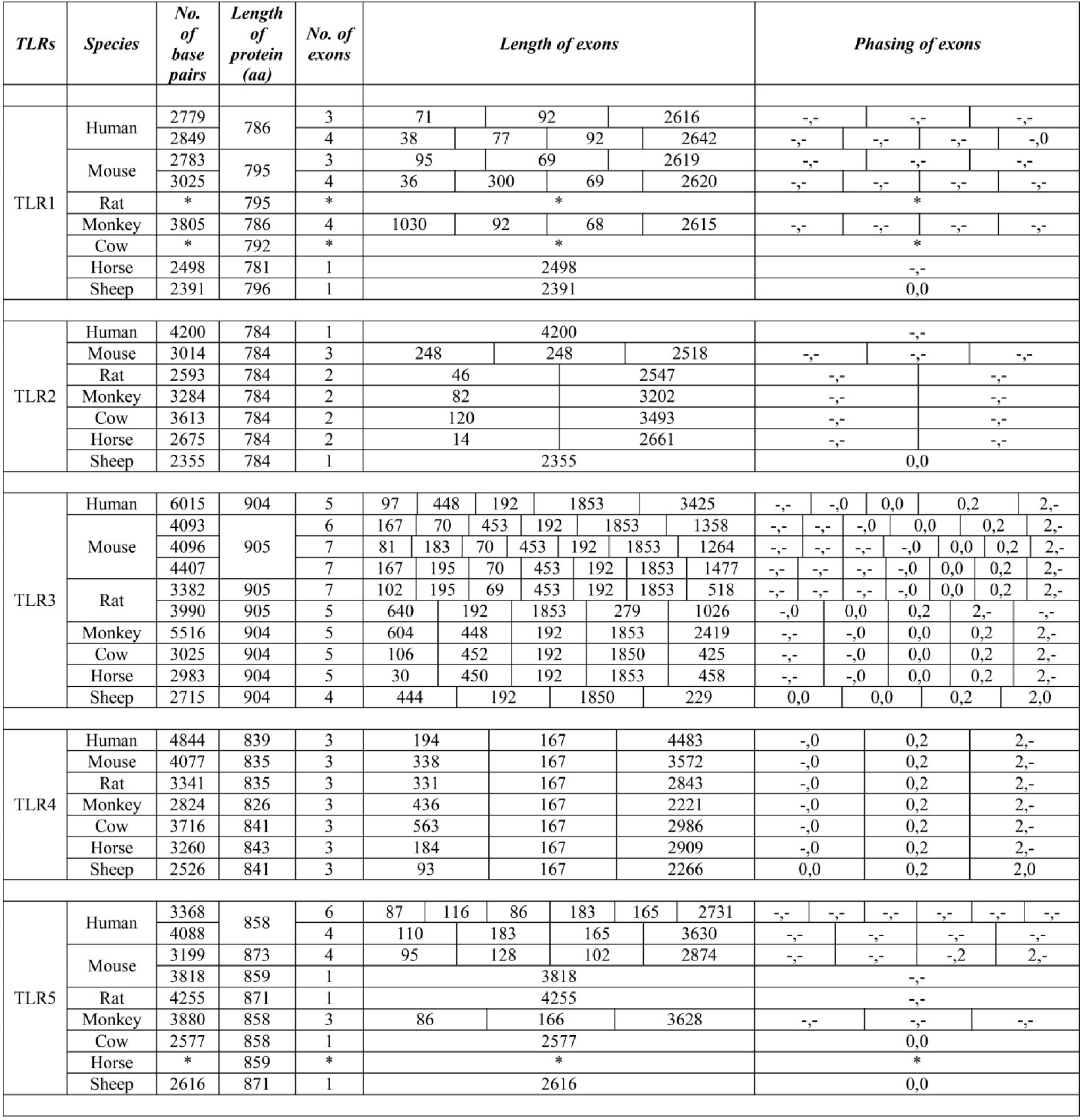
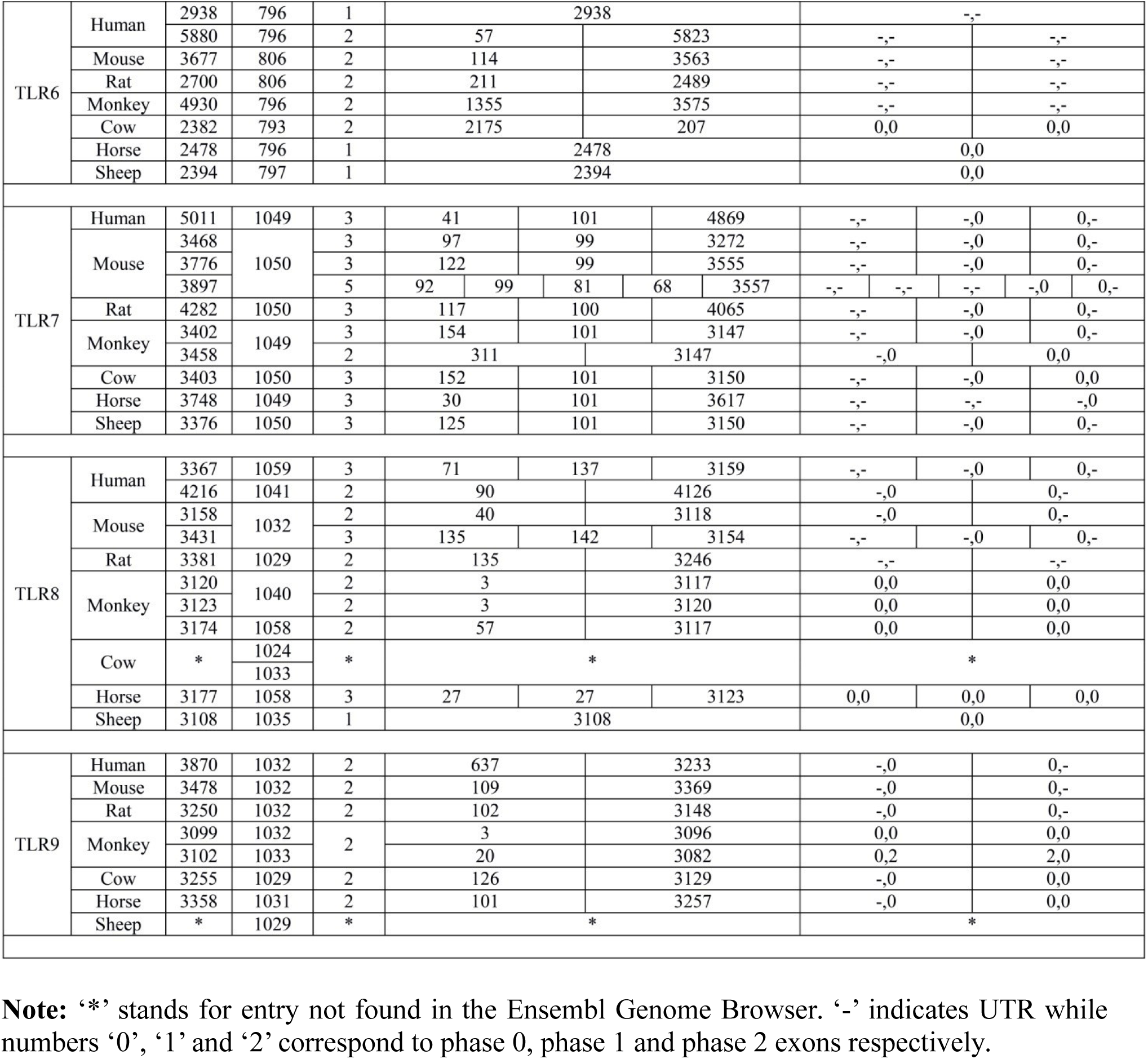
Chart showing base pair length (bp), length of proteins, number, length and phasing of exons in the nine TLRs across seven species.

The number of exons identified within each TLR ranges from a minimum of 1 to 7. There has been a difference in the number of exons across seven organisms even within a specific TLR except for TLR4 and TLR9. TLR4 and TLR9 consist of 3 and 2 exons, respectively, for all the metazoans. Barring TLR6 and TLR7, the other TLRs (TLR1, 2, 3, 5 and 8) show considerable variation in the number of exons. Interestingly, codon interruption seems to be a rare event in the TLR family with TLR3 and TLR4 as exceptions. In this case, disruption of codon occurs at the same position in all the species. Also, codon disruption has been noticed in one of the transcript variants of TLR5 and TLR9 for a single organism. Another similarity observed between TLR3 and TLR4 is that both possess an exon of invariant length for all species, a feature unseen in other members of the TLR family. Unlike TLR3 (4, 5, 6 & 7 exons), the position of the exon in TLR4 is restricted owing to the fixed number of exons (3 exons) in all the species.

We concentrated on TLR4 further in the gene study as we observed a high degree of conservation in the number and phasing pattern of exons. Out of the three exons encoding TLR4, the second exon possesses a strikingly conserved CDS of 167 base pairs across all the seven organisms (Figure 2). Two different multiple sequence alignments were carried out with the second exon and with its corresponding amino acid sequence (Figure 3a and 3b).

**Figure 3a.**
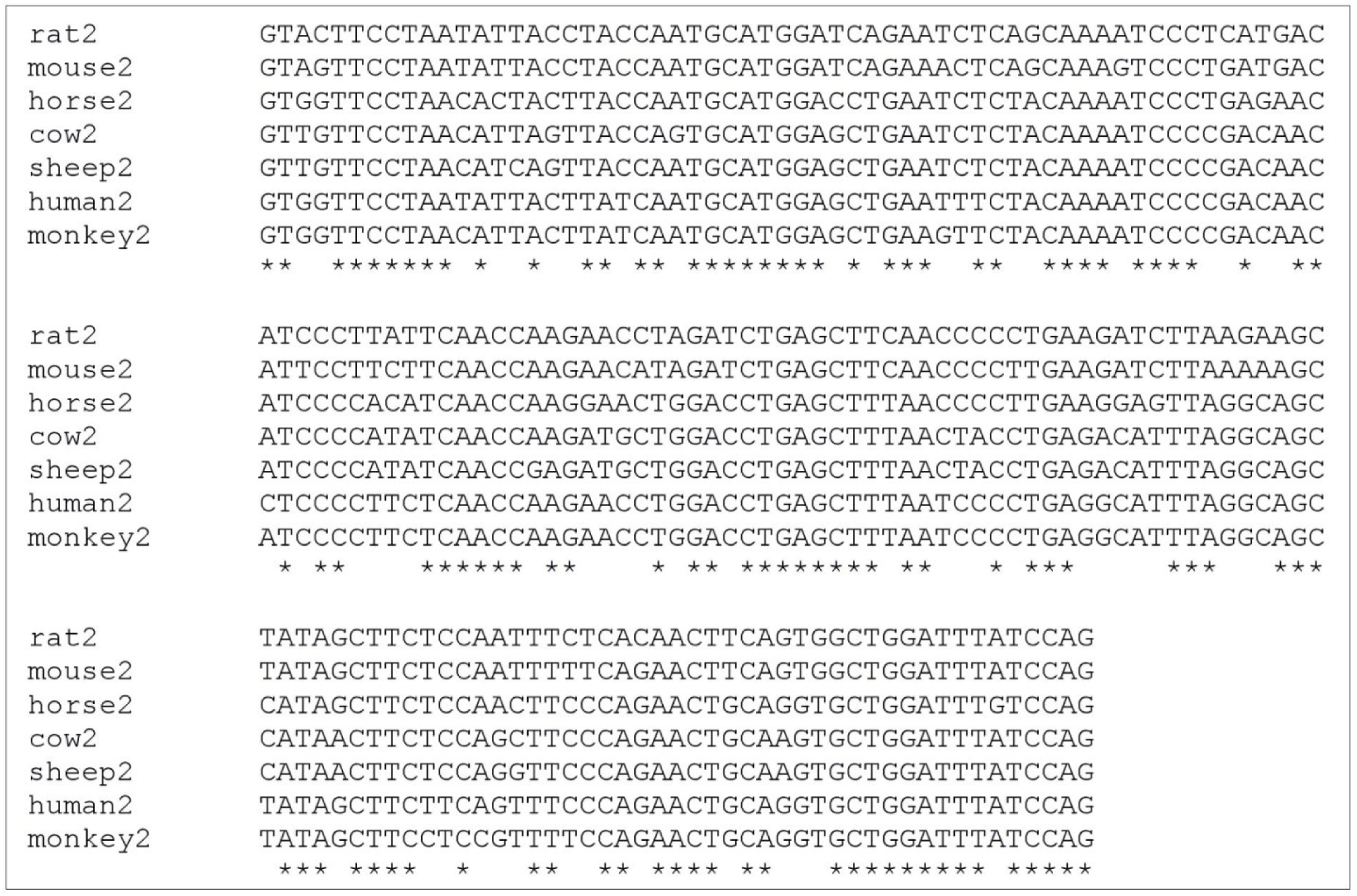
Multiple sequence alignment of the coding sequence corresponding to the second exon of TLR4 from seven metazoans.

**Figure 3b.**
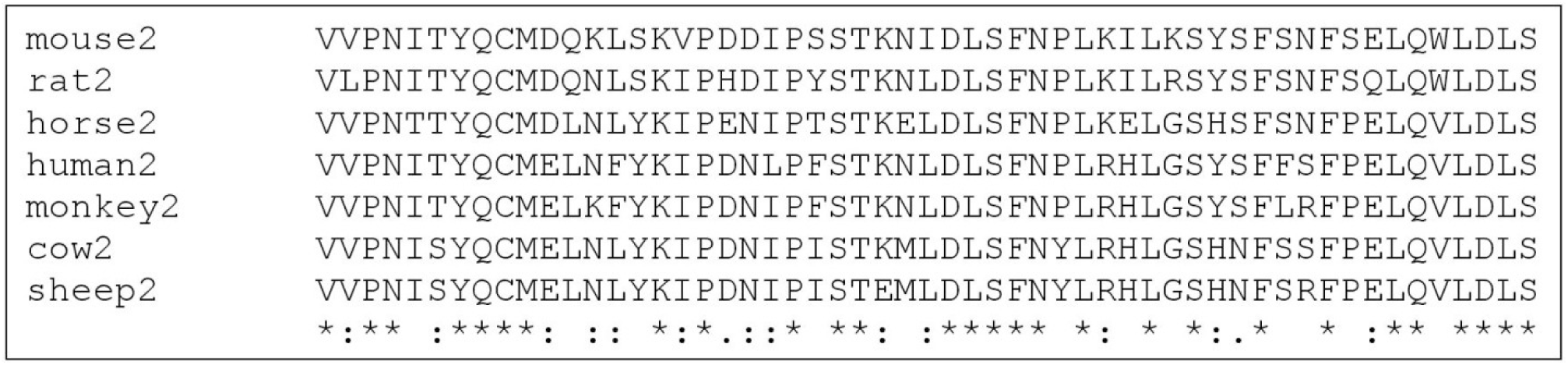
Multiple sequence alignment of the amino acid residues encoded by the second exon of TLR4 from seven metazoans.

The multiple sequence alignments of exon and amino acids confirmed that the conserved pattern which prevails in the second exon of TLR4 is retained after translation. We can possibly infer that there is a selective constraint in maintaining the length as well as the nature of residues in this region. A very pertinent question that arises at this point is that whether the second exon is coding for LRRs, if so then how many of them and which part of the LRR (HCS or VS) is being coded that demands such a level of conservation.

### 3.3 LRRs within TLR4

As mentioned above, the unique organization of the gene structure of TLR4, exclusively within the metazoans, lead us to work with this member of TLR. Primarily, we identified 23 tandem leucine-rich repeats (LRR1, LRR2, LRR3……LRR23) among nine different organisms by manual annotation of multiple sequence alignment (Figure 4) considering the conserved pattern to be L x x L x L x x N/C x L where L= Leucine (L), Isoleucine (I), Valine (V) and Phenylalanine (F), N= Asparagine (N), Cysteine (C), Serine (S) and Threonine (T). It is to be noted that similar alignment was also obtained from T-Coffee (M-Coffee) (Supplementary Figure S1). Length of 23 LRR motifs, identified manually in TLR4, ranges from 21 to 31 residues with their HCS fixed to 11 residues and VS varying from 10 to 20 residues. Few LRRs such as LRR3, 4, 6, 7, 15, 16, 17, 20 and 23 follow the typical pattern for LRRs without any substitution or conserved mutations in place of ‘L’ in the 1^st^, 4^th^, 6^th^ or 11^th^ position. In case of LRR1, we observe conserved methionine substitutions in the 9^th^ position in place of ‘N’ whereas the same is evident in the 6^th^ position in place of ‘L’ for both LRR19 and LRR21. Substitutions of ‘L’ with ‘T’ are seen in LRR2 (in all organisms at the 1^st^ position), LRR12 (in 4 organisms at the 1^st^ position), LRR18 (in 5 organisms at the 11^th^ position) and LRR22 (in 2 organisms at the 6^th^ position). In LRR10 and LRR11 substitution of ‘N’ at the 9^th^ position with ‘L’ and ‘F’ are observed, respectively. In LRR1 and LRR22, ‘Y’ is seen to occupy the 6^th^ position (in all organisms) and 9^th^ position (for 2 organisms). In few LRRs such as in LRR8, 9 and 14, we observe ‘N’ and ‘D’ in place of ‘L’ in many organisms. We also notice, ‘L’ changing to ‘A’ and ‘W’ in LRR5 and LRR13 respectively for all organisms. Moreover, substitutions of ‘L’ with hydrophilic amino acid residues are observed at many positions in the HCS of LRR motifs. Also, conserved residues in the otherwise variable ‘x’ positions of the HCS of LRRs across nine organisms may imply structural and functional importance.

**Figure 4.**
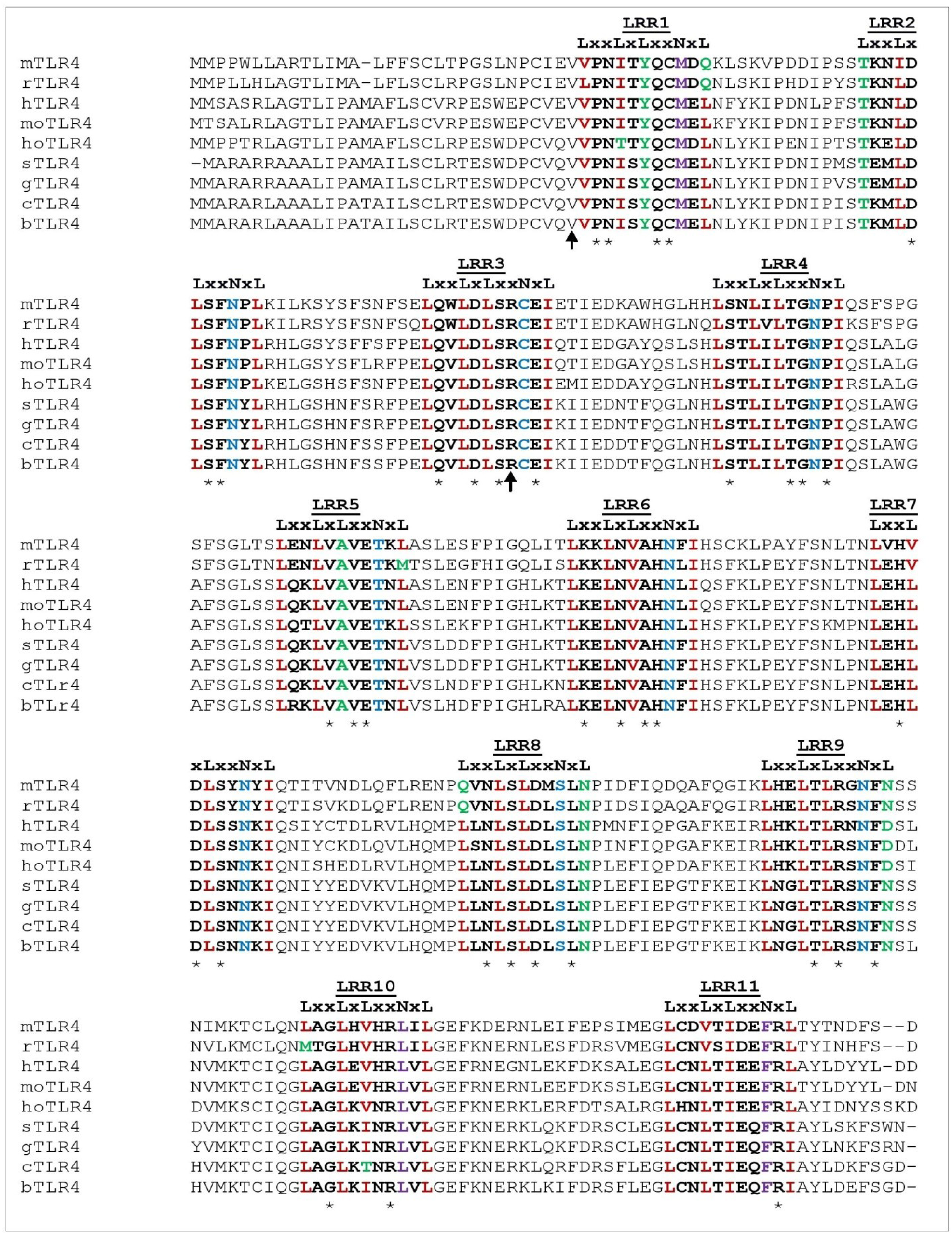
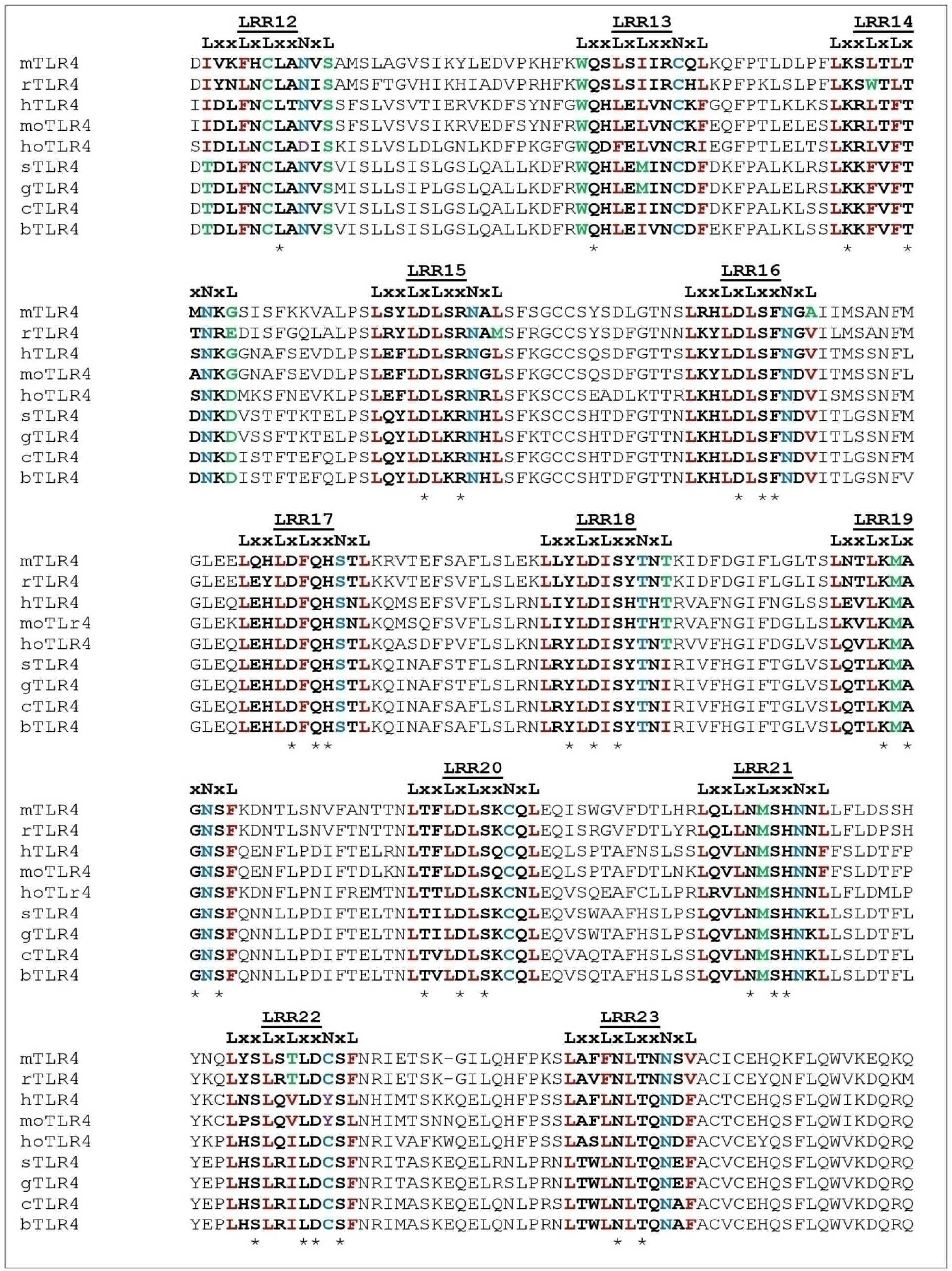
Multiple sequence alignment of the ectodomain within TLR4 among nine metazoans. **Note:** The number of LRRs in the extracellular domain does not include the LRR-NT and LRR-CT motifs. The organisms used for MSA are abbreviated as follows-mouse (mTLR4), rat (rTLR4), human (hTLR4), monkey (moTLR4), horse (hoTLR4), sheep (sTLR4), goat (gTLR4), cow (cTLR4) and buffalo (bTLR4). The conserved part of the LRR motifs (LxxLxLxxNxL) in the entire MSA have been marked with a user defined color code where positions 1, 4, 6 and 11 containing either leucine (L), isoleucine (I), valine (V) or phenylalanine (F) are colored in red, whereas any other substitution taking place in either of these positions are coded in green. Similarly, position 9 in the conserved segment consisting of asparagine (N), threonine (T), cysteine (C) or serine (S) is marked in blue and any change in this position is coded in purple. Highly conserved residues in the ‘x’ position of all organisms has been marked with ‘*’. The two arrow signs indicate the start of the second and the third exon respectively.

Trying to map the LRRs of TLR4 with respect to the encoding exons (3), it may be pointed out that the first exon is devoid of any LRR and the second exon consists of LRR1, LRR2 and a portion of LRR3 with the rest of the LRRs, TM and the TIR domain residing entirely in the third exon. Furthermore, we notice the interacting residues of TLR4, with MD-2 (dimerizing partner) and LPS (ligand) are spread over LRR10, 11, 13, 14, 15, 16, 17 and 18 (Park et al., 2009). Despite the fact that the ligand interacting residues lie in LRRs far away from LRR1, LRR2 and portion of LRR3, what calls for such a level of stringency at the gene and protein level, remains unanswered. A schematic diagram showing the domain architecture of TLR4 has been shown in Figure 5.

**Figure 5.**
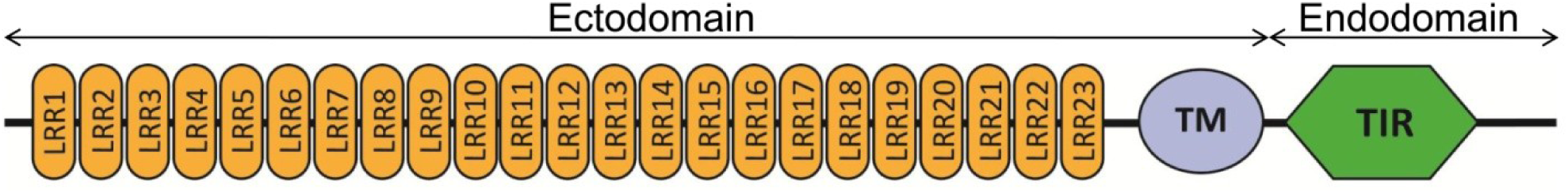
Schematic representation of the domain architecture of TLR4.

### 3.4 Phylogenetic tree analysis

#### 3.4.1 Human TLRs

An unrooted phylogenetic tree was constructed with 1000 bootstrap values using parsimony method for nine TLR protein sequences of human (Figure 6). It may be mentioned that similar tree (Figure 7) was obtained from the sequences without TIR domain (ectodomain consisting of LRRs and TM).

**Figure 6.**
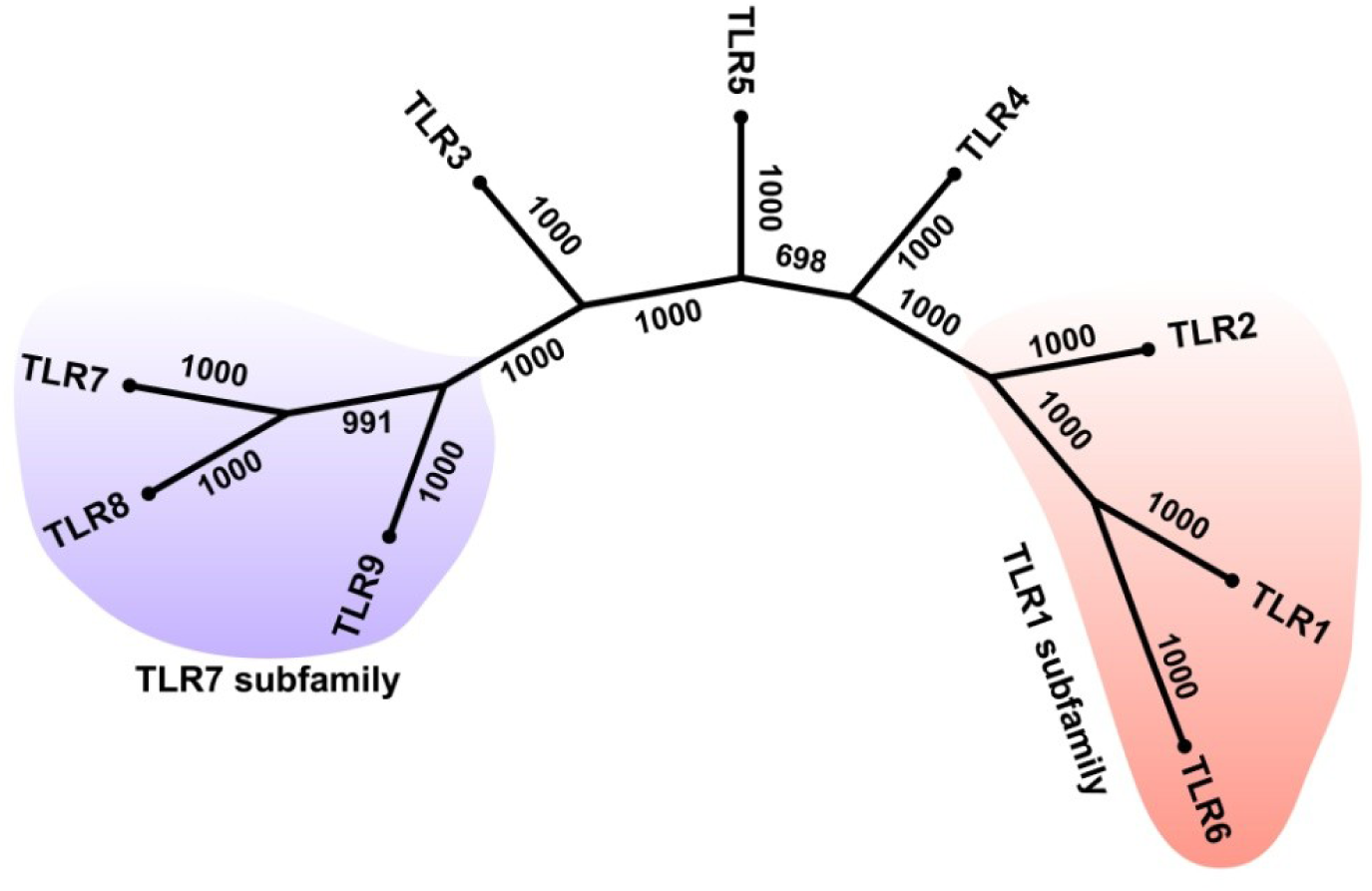
Phylogenetic tree representing the whole protein sequences of nine human TLRs (TLR1-TLR9)

**Figure 7.**
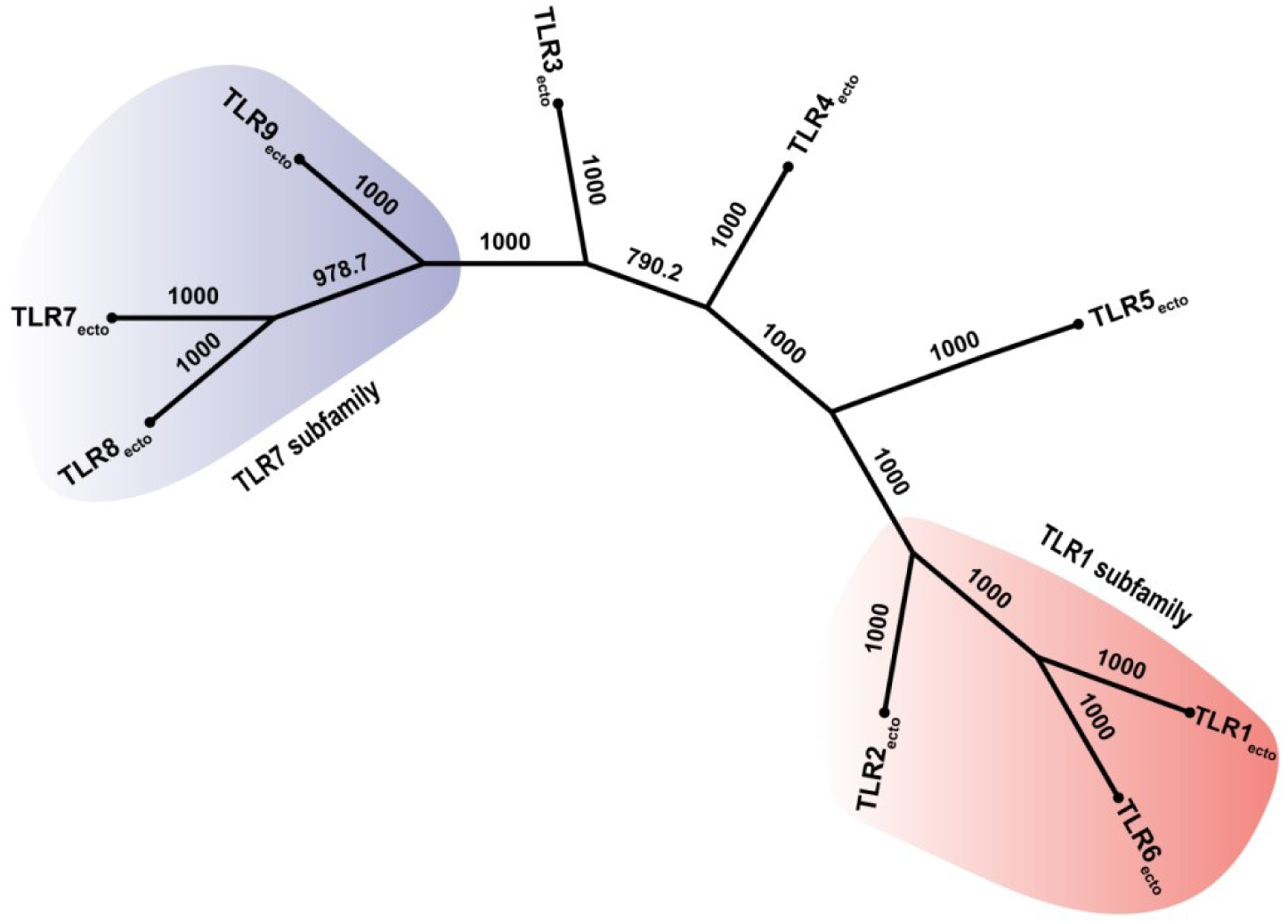
Phylogenetic tree of the ectodomain of the nine human TLRs.

TLRs (TLR1, TLR2 and TLR6) with comparatively shorter length form a cluster while the longer TLRs (TLR7, TLR8 and TLR9) form another. The topologies of the two trees (whole protein and ectodomain) reflect their functional similarity in terms of ligand recognition. The TLRs that identify bacterial lipopeptides (TLR1, TLR2 and TLR6) constitute one group and those that recognize viral or microbial nucleic acids (TLR7, TLR8 and TLR9) seggregate to another. Although TLR3 share similarity with TLR7, TLR8 and TLR9 with respect to recognition of viral nucleic acids, yet it is relatively of shorter length compared to the other three. In fact, this justifies its distinct position in the phylogeny. Also, TLR4 and TLR5 occupy a distinct position in the tree without clustering in any of the TLR families. Besides, TLR4 and TLR5 are of almost equivalent length and recognize ligands (lipopolysaccharides and flagellin, respectively) that are of completely different nature from the other TLRs. Figure 6 and 7 show different line of evolution for TLR3, TLR4 and TLR5. It may be suggested that determination of domain architecture and phylogenetic analysis restates LRR-containing TLRs to have evolved through independent evolution of the ectodomain.

A rooted phylogenetic tree was contrived for the TIR domain of nine human TLRs and TIR domain of SARM1 (Sterile alpha and TIR motif-containing protein 1). SARMs are the only known TIR-containing adapters that negatively regulate TLR signalling and have a different line of evolution from other TIR-containing adapters. Therefore, TIR of SARM1 was treated as an outgroup to establish the clustering pattern of TLRs in the presence of a non-TLR TIR domain. Despite the similarities in the length of both TLR and non-TLR TIR domains, SARM1 stands out as an operational taxonomic unit in the tree. Since the TIR domain is evolutionarily conserved, we wanted to see the origin of the nine TIRs, the only part being least susceptible to changes by selection pressure. From this tree (Figure 8), it is clear that TIR domains of TLR3 and TLR5 might have undergone extensive divergence in comparison to the other members. The fact that TIR of TLR4 shares some similar adapter molecules for signal transduction with TLR2 (Akira and Takeda, 2004; O’Neill and Bowie, 2007), justifies the position of TLR4 in the tree.

**Figure 8.**
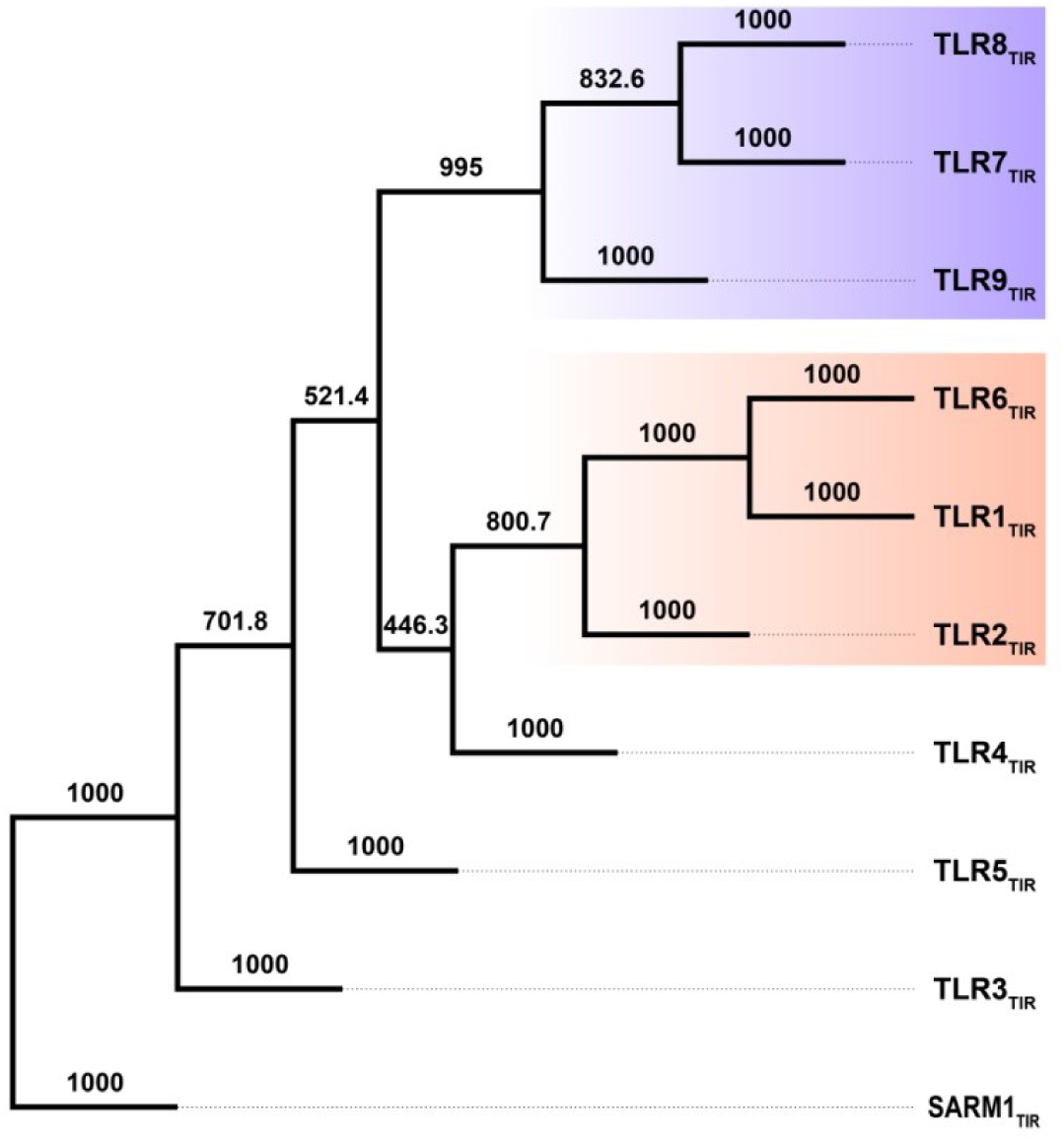
Phylogeny considering the TIR domain of the nine human TLRs.

#### 3.4.2 TLR4

We have drawn separate unrooted phylogenetic trees taking both protein and gene sequences of TLR4 from seven organisms (Supplementary Figure S2a and S2b). Analyses exhibit identical clustering pattern at the gene as well as the protein level implying significant conservation. Since major part of these proteins comprise of repeats that are only LRRs, it would be intriguing to find out if the repeats at different positions from the same organism or repeats from different organisms are invariant or distinct in nature. Consequently, phylogenetic trees considering the amino acid and coding sequences of portions corresponding to 23 LRR motifs in TLR4 across seven organisms, each with 1000 bootstrap iterations, were crafted (Figure 9a and 9b). Thus, a total of 161 (23LRRs × 7organisms) sequences were taken into account for constructing each of these unrooted trees. It is worth mentioning that the 23 repeats from seven organisms form distinct clades both at the protein and the gene level indicating unique spatial arrangement of the repeats. At the organism level, evolution of the repeats maintaining their individuality, points towards structural and functional implications. It may be mentioned that some LRR pairs in the tree of protein sequences appear to be proximate under a common node. Such LRR pairs are LRR2-LRR16, LRR3-LRR20, LRR4-LRR18, LRR6-LRR19, LRR10-LRR23 and LRR12-LRR22. While in the gene tree, the pairs that are arranged under a common node are LRR1-LRR17, LRR2-LRR22, LRR6-LRR19, LRR8-LRR16, LRR10-LRR13, LRR11-LRR12 and LRR15-LRR23. To sum up it may be inferred that duplication event might have given rise to the repeats mentioned above leaving out LRR5, LRR7, LRR9, LRR14 and LRR21 to have evolved independently.

**Figure 9a.**
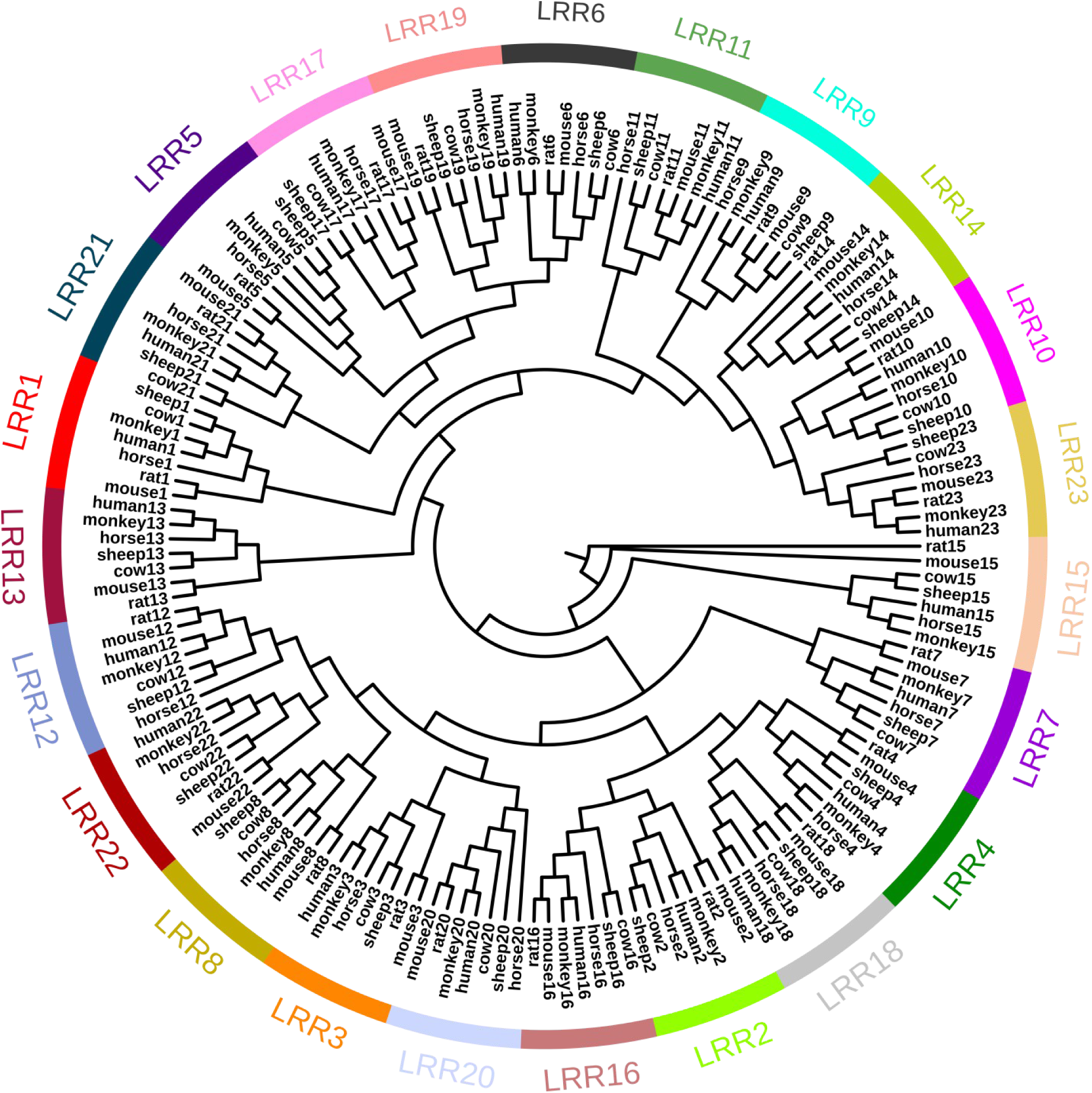
Phylogenetic tree with the protein sequence of 23 individual LRR motifs identified in TLR4 across seven organisms.

**Figure 9b.**
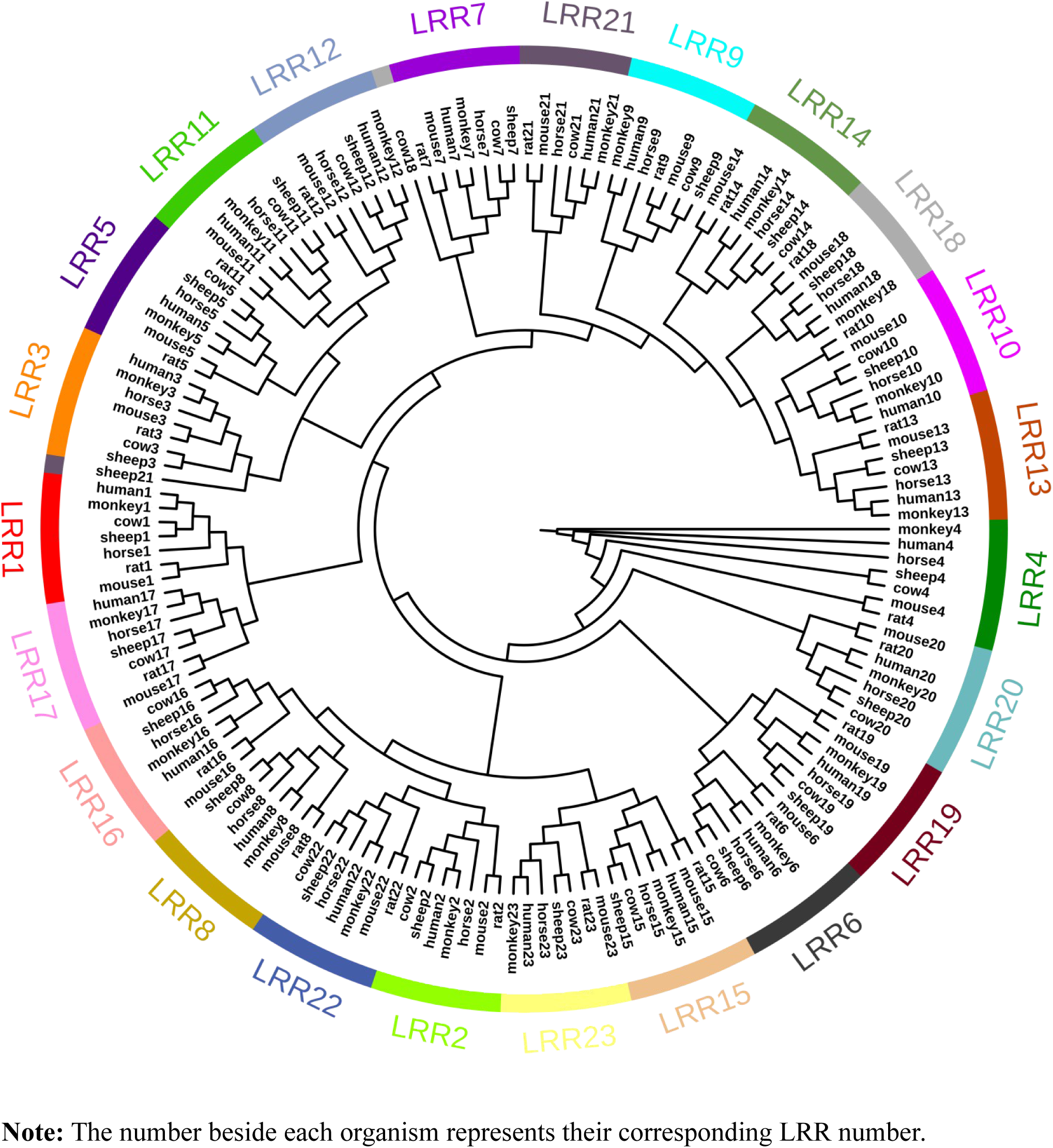
Phylogenetic analysis with the coding sequence of 23 individual LRR motifs across seven organisms.

### 3.5 Selection pressure on individual LRR coding sequences

Selection pressure on genes can be identified from the ratio of non-synonymous (K_a_) to synonymous (K_s_) nucleotide substitution with pairwise or multiple combinations of genes. We have calculated K_a_ and K_s_ of 23 LRR coding sequences from seven organisms using the calculation tool mentioned earlier. Coding sequences corresponding to 23 repeats were translated to protein sequences, multiply aligned on codon boundaries to construct phylogenetic tree with K_a_ and K_s_ values. Figure 10 shows the differential selection pressure on branches of gene phylogenetic tree for LRR11, LRR14, LRR6 and LRR21. A ratio significantly greater than one indicates positive selection pressure which has been found in almost all LRRs of TLR4 (Supplementary Table S1). LRR11 and LRR14 needs special mention as they exhibit strong positive selection for most of the organisms considered. The probable reason may be attributed to the fact that the residues responsible for recognition of its ligand and MD-2 (Park et al., 2009), lies in 11^th^ and 14^th^ repeat. Additionally, it may be mentioned that more selection pressure acts on these two LRRs particularly to impart a wide range of recognition ability for diverse array of bacterial LPS. On the other hand, LRR6 and LRR21, known not to directly participate in recognition, are seen to undergo purifying selection.

**Figure 10.**
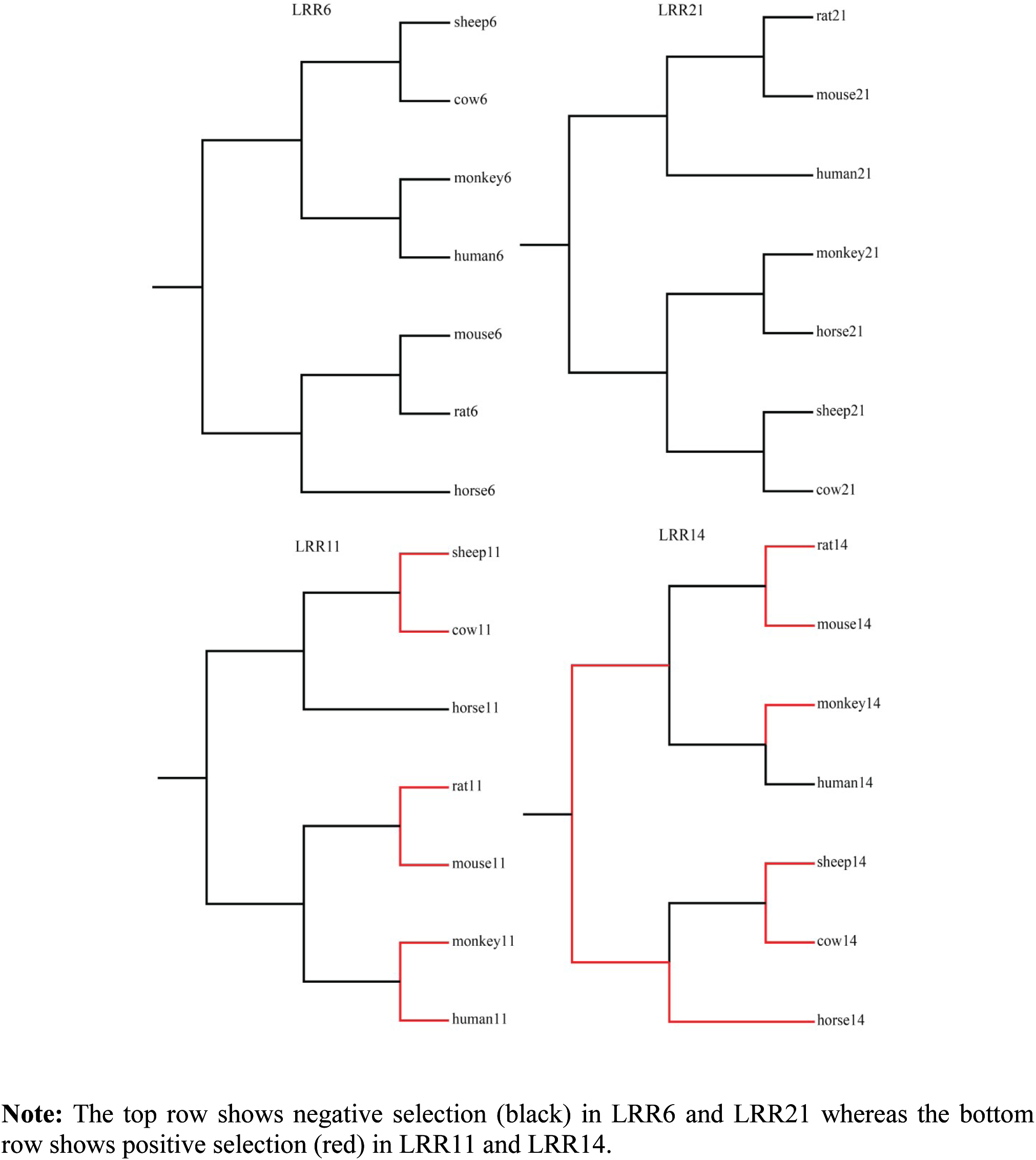
K_a_/K_s_ ratio tree showing selection pressure in four particular LRRs.

### 3.6 Positive selection in the ectodomain of TLR4

According to our results obtained from the previous section, positive selection prevails on specific LRR motifs in the ectodomain part of TLR4 across seven organisms. The selection pressure is constrained to particular repeats of functional importance instead of being distributed over the entire ectodomain region, in other words spanning the repeat motifs. It would thus be intriguing to verify if this specificity in sites for selective pressure is a general rule applicable to other TLRs or not. Consequently, we performed pairwise codon alignment between human and monkey using PAL2NAL considering the ectodomain, endodomain and entire sequence of nine TLRs (Figure 11). From the results, we observe that the value decreases considerably for the viral or bacterial nucleic acid sensing TLRs (i.e. TLR3, 7, 8 & 9). Our result is shown to be consistent with the findings that evolution in the aforementioned TLRs is constrained thus avoiding response against self-derived nucleic acids (Krieg and Vollmer, 2007; Wlasiuk and Nachman, 2010). The same argument does not hold true for the low selection rate in TLR6 ectodomain as it does not belong to the above mentioned TLR family. TLR4 shows highest selection rate among the other TLRs thereby establishing as a special TLR in terms of exon phasing pattern, recruitment of adapter molecules in alternate basis for signalling and recognition of a myriad of LPS ligands.

**Figure 11.**
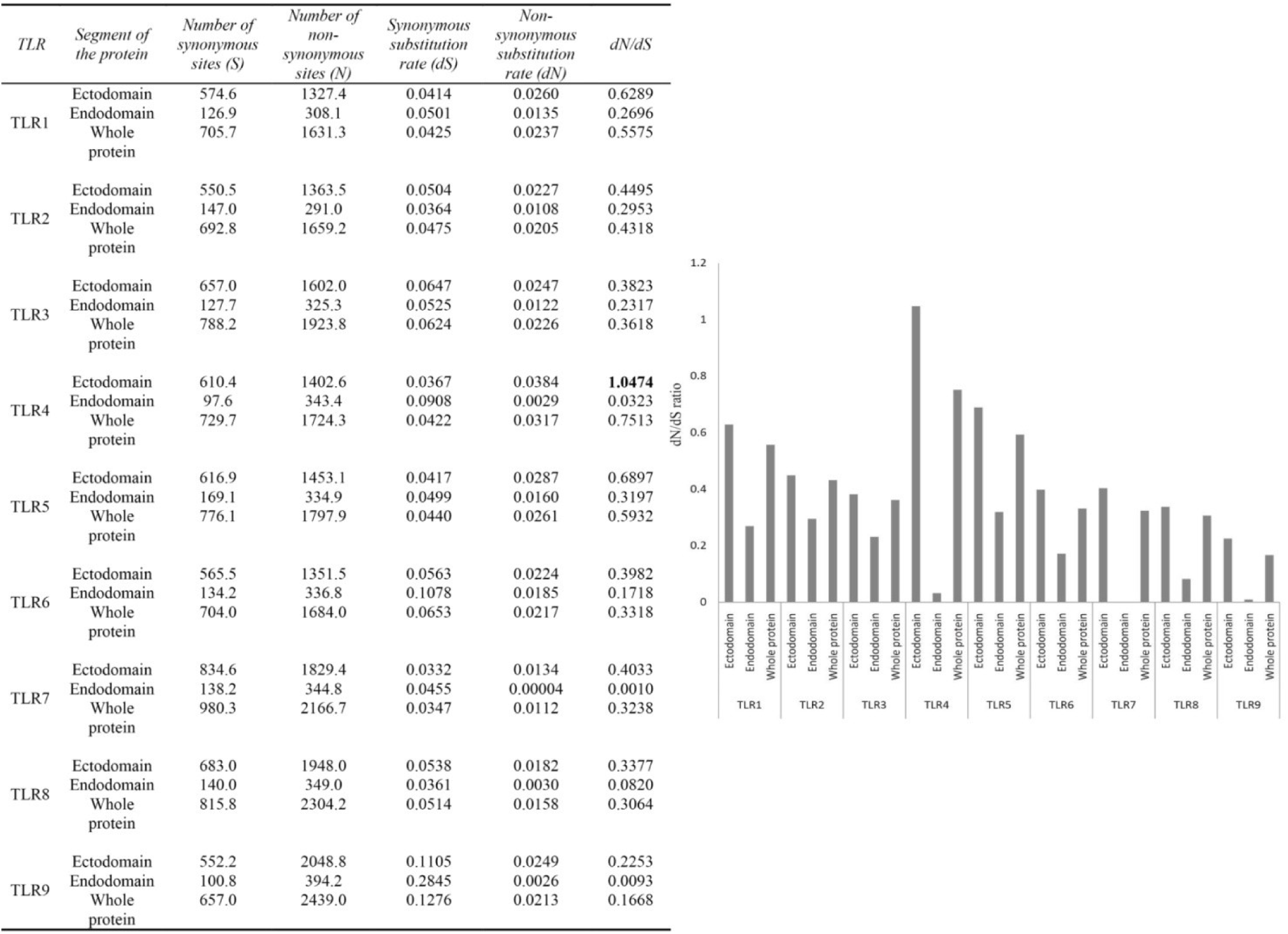
Tabular and graphical representation of the dN/dS calculations between human and monkey in nine TLRs.

## Conclusion

Domain architecture analysis of TLRs revealed that each TLR consists of numerous repeat-spanning ectodomain, a transmembrane region and a single TIR domain. On verification of the repeat number using different databases, it was found that the LRR motifs are present in distinct numbers in each TLR remaining invariant across species. Gene structure of TLRs revealed that the number of exons encoding the proteins varied across species for the same TLR and also across different TLRs. TLR4 and TLR9 is an exception to this as both are encoded by 3 and 2 exons respectively across all species studied. Unlike the coding pattern of ribonuclease inhibitor an LRR protein, the repeats of TLR are not coded by individual exons. For example, there are cases where few repeats have been found to be coded by a single exon as is a single repeat found to be coded by two exons. Interestingly, only two types of phasing of introns, phase 0 and phase 2 were observed. Phase one introns were missing from all members of the TLRs studied here. Codon interruption has been a rare event in these proteins as introns with phase 2 was observed for only TLR3 and TLR4 thus supporting the early intron hypothesis for this family of proteins. It is worthy of note that the second exon of TLR4 remains conserved both at the CDS and its translated level. According to MSA results, the length and sequence residues of this portion are conserved across all the metazoans studied. Despite being devoid of any ligand or adapter interacting residues in this region, the aforementioned conservation may probably have either some structural implications or functional relevance. Further study of TLR4 ectodomain at the protein level indicated that 23 distinct repeats, following the typical LRR signature sequence, are present in tandem array across seven organisms. Phylogenetic analysis, considering different segments (ectodomain, endodomain and whole protein) of TLR1-9 separately, revealed that TLR3, TLR4 and TLR5 occupy distinct positions between the TLR1 and TLR7 subfamilies. Phylogeny at the CDS and protein level, with individual repeats of the TLR4 ectodomain, reported the presence of each LRR as a separate clade for all the organisms, supporting the fact that individual repeats have undergone distinct evolution. On analysis of selection pressure, we found that the K_a_ and K_s_ values vary differentially among organisms for each motif considered. Out of all, two repeats, LRR11 and LRR14, exhibited strong positive selection on most of the organisms while two others, LRR6 and LRR21, exerted negative selection pressure. Both LRR11 and LRR14 harbour ligand and adapter interacting residues thus explaining the cause for positive selection. The cause of negative selection for LRR6 and LRR21 could be for structural maintenance of the scaffold for efficient recognition. Evaluation of selection pressure on different segments of these PRRs (pathogen recognition receptor) revealed a skewed selection pressure along the length of TLR4. The ectodomain consisting of the repeats exhibited positive selection while the endodomain or the TIR domain showed negative selection. The positive selection of the ectodomain is in accordance with its demand of interacting with the ever changing PAMPs to facilitate efficient recognition eventually activating the signalling cascade through the TIR domain conserved across the TLR family.

## Acknowledgements

The authors are thankful to use the facilities provided by the Centre for High-Performance Computing for Modern Biology (CHPC) in the University of Calcutta. We express our sincere gratitude to Dr. Aditi Maulik, Postdoctoral fellow at the Indian Institute of Science (IISc) and Sucharita Das, Senior Research fellow at the University of Calcutta, who provided insight and expertise that greatly assisted this research work.

