## Supplementary Materials for "Insights into the evolution of extracellular leucine-rich repeats in metazoans with special reference to Toll-like receptor 4"

**Figure S1** Multiple sequence alignment of the ectodomain within TLR4 using M-Coffee

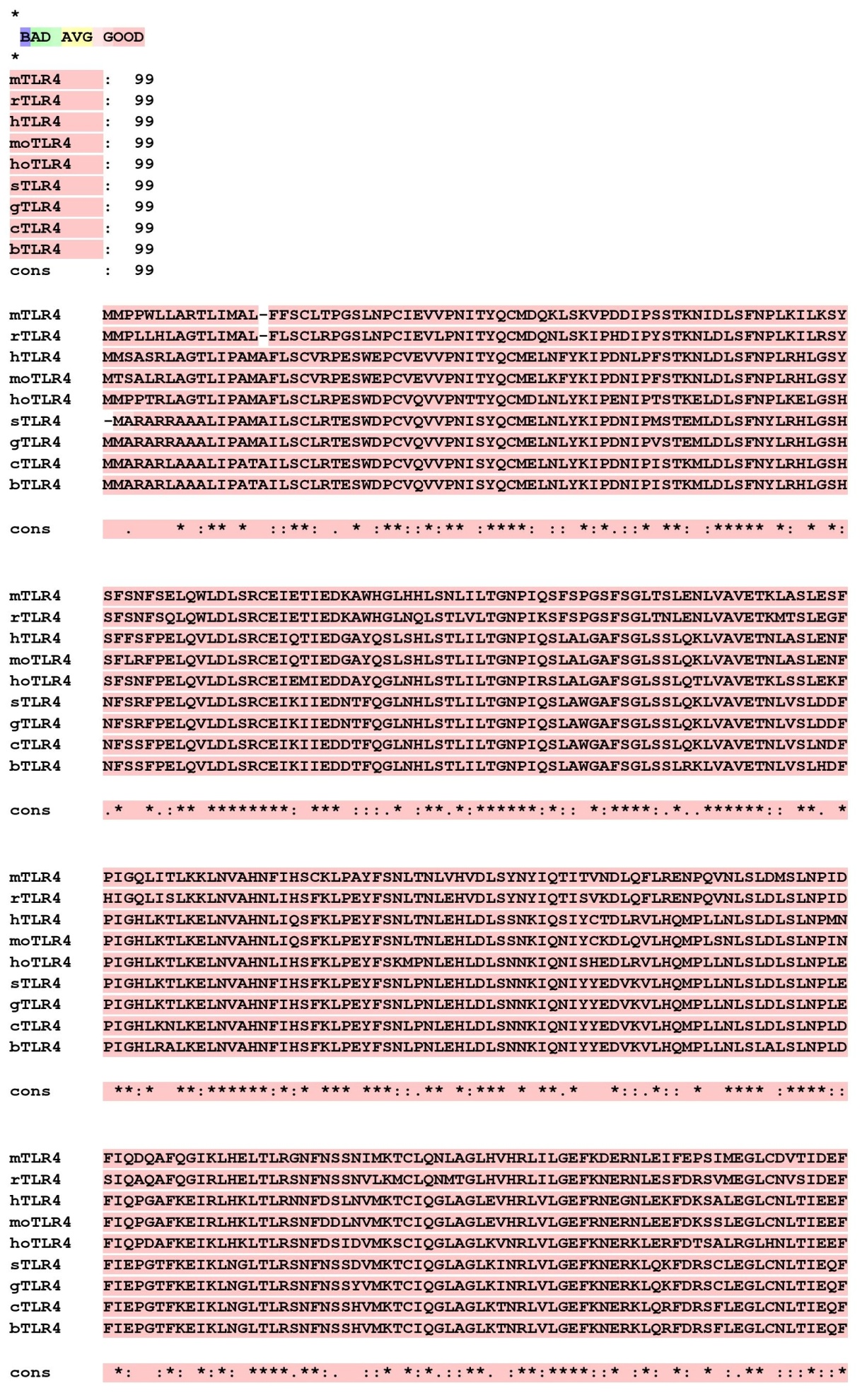

**Figure S1 (continued)**

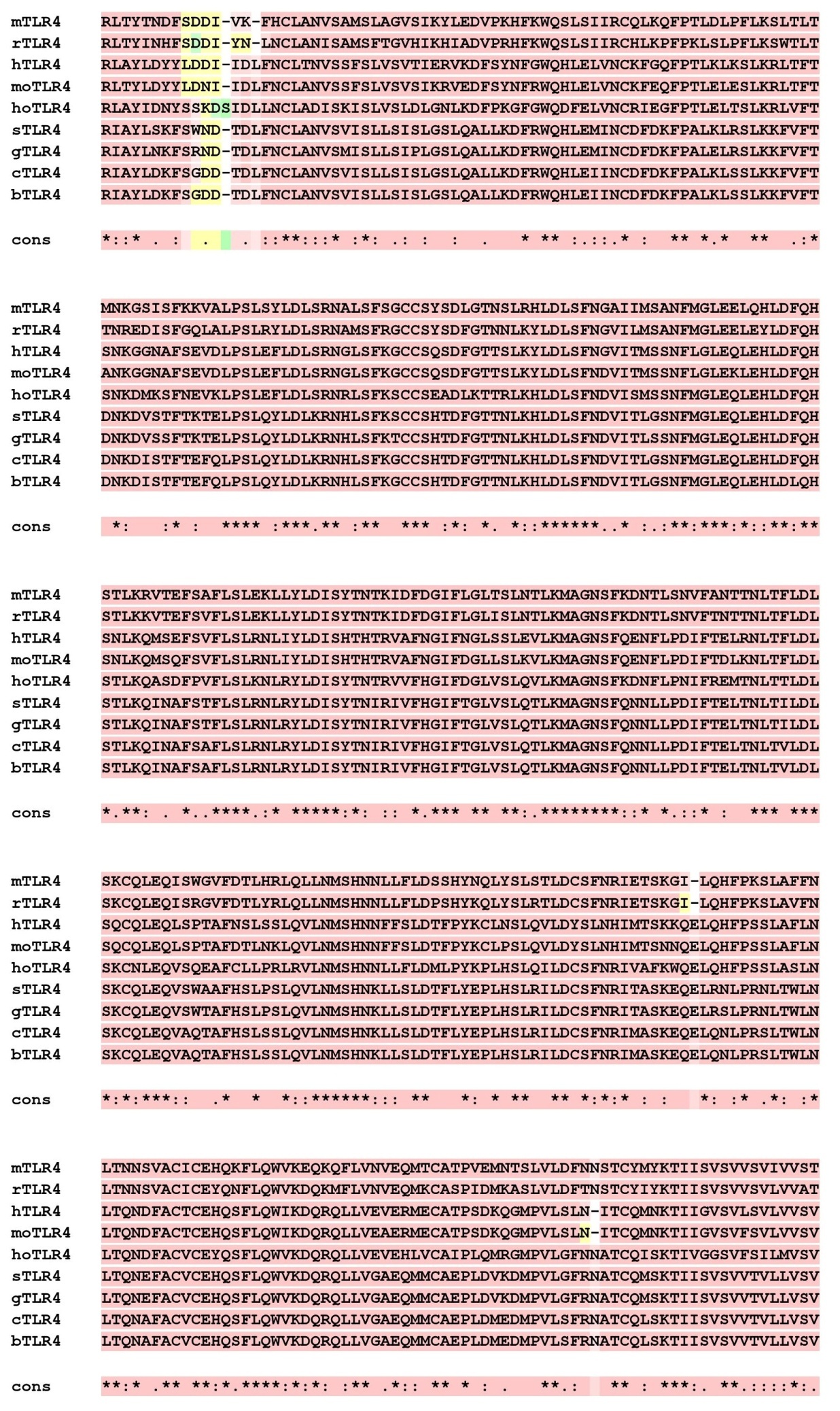

**Figure S2a** Phylogenetic tree of the TLR4 protein across seven metazoans

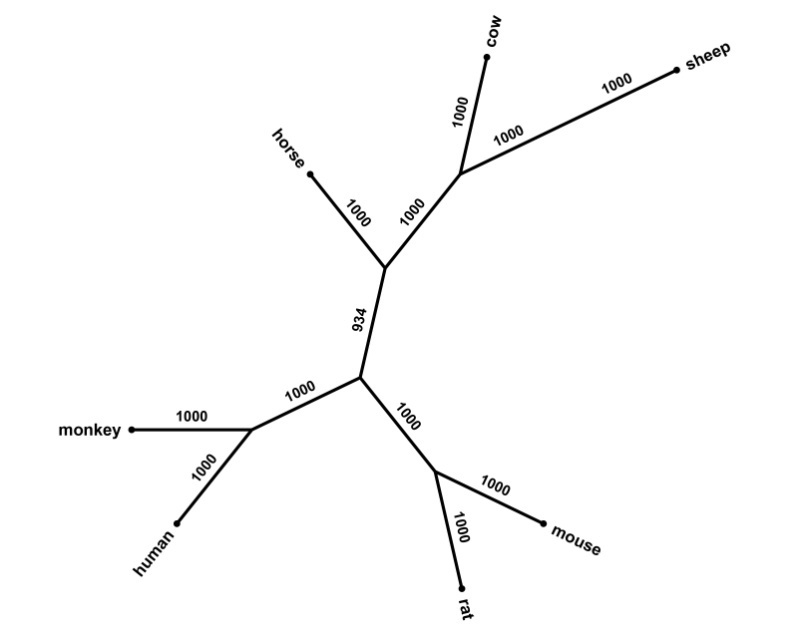

**Figure S2b** Phylogenetic tree of the TLR4 coding sequence of seven metazoans

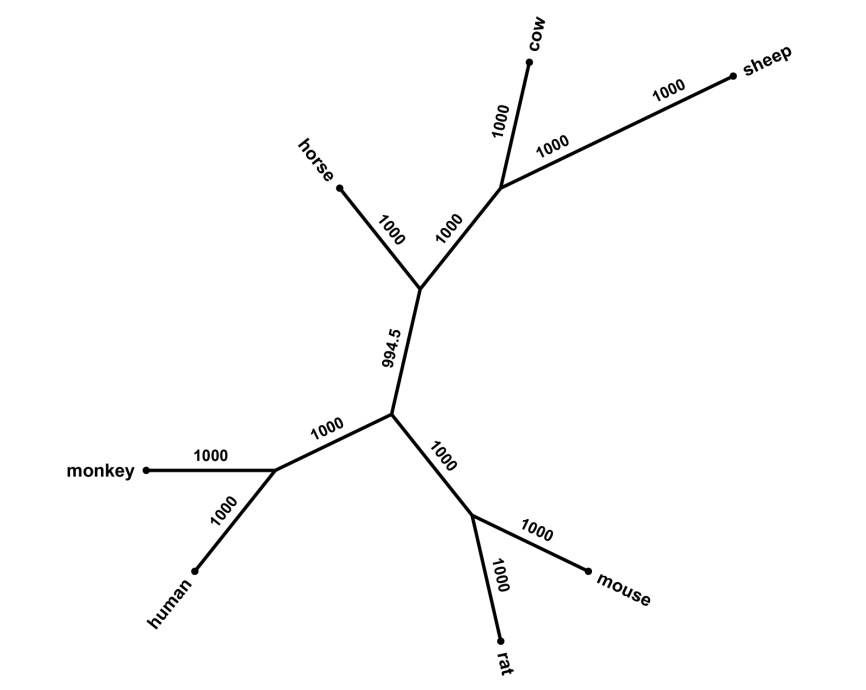

**Table S1** showing the K_a_/K_s_ ratio of 23 LRRs within TLR4 among seven organisms

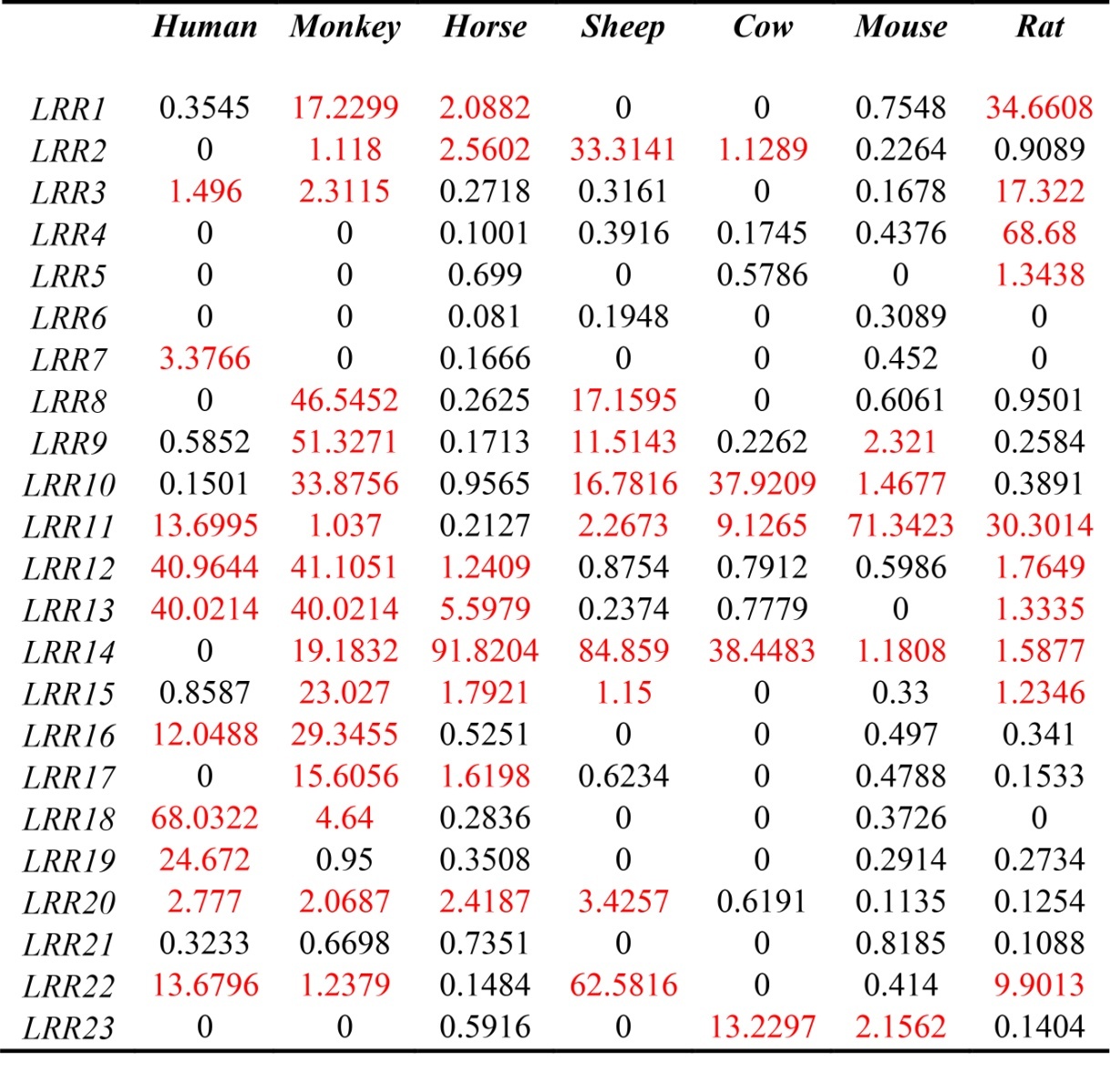
